# Pair correlation function analysis revealed different nuclear translocation mechanisms for glucocorticoid receptor’s monomeric and dimeric forms

**DOI:** 10.64898/2026.06.05.730430

**Authors:** María Cecilia De Rossi, Diego Martín Presman, Valeria Levi

## Abstract

Glucocorticoids are among the most widely prescribed drugs globally due to their potent anti-inflammatory and immunosuppressive actions. These effects are primarily mediated by the glucocorticoid receptor (GR), a ligand-activated transcription factor that translocates from the cytoplasm to the nucleus to regulate hundreds of genes. Although nuclear entry is a prerequisite for its genomic response, the mechanisms governing this process remain unresolved; specifically, whether the receptor translocates as a monomer or a dimer remains a subject of significant controversy. Here, we employed the pair correlation function (pCF) approach to quantify the nuclear translocation of single fluorescent GR molecules in live cells. This minimally invasive method identifies correlations between intensity fluctuations generated by molecules moving from the cytoplasm into the nucleus. Our results demonstrate that GR’s quaternary structure and conformation modulate GR transport. While GR monomers rely exclusively on passive diffusion, GR dimers also utilize the microtubule-dynein machinery for active transport, proving that dimerization can precede nuclear import. Furthermore, the perinuclear vimentin network facilitates faster translocation by constraining actively transported dimers near nuclear pores. Collectively, our work reconciles contradicting reports regarding GR stoichiometry during import by demonstrating that both monomers and dimers translocate, albeit through distinct mechanisms. Importantly, these results reopen the door for a microtubule-dependent, heterocomplex-independent model of GR translocation, suggesting that the cytoskeleton is an integral, yet overlooked, component of the GR signaling pathway.

## Introduction

Glucocorticoids are steroid hormones that act as critical physiological regulators by controlling metabolism, stress responses, cardiovascular function, and many aspects of development and reproduction [1]. This broad influence makes them indispensable in both health and disease. In fact, they are among the most widely prescribed drugs worldwide due to their potent anti-inflammatory and immunosuppressive action [2]. However, their chronic use can lead to severe side effects, including osteoporosis, metabolic disorders, and an increased risk of cardiovascular disease [3].

Glucocorticoids act primarily by binding and activating the glucocorticoid receptor (GR), a transcription factor (TF) expressed in nearly all human cell types. GR regulates a diverse set of gene networks in a context-dependent manner, explaining the widespread effects of glucocorticoids in both physiology and disease [4].

Structurally, GR shares the characteristic modular organization of steroid receptors, comprising a central DNA-binding domain (DBD), a C-terminal ligand-binding domain (LBD) separated from the DBD by a flexible *hinge* region, and two activation function regions (AF1 and AF2) located in the N-terminal domain (NTD) and LBD, respectively [5]. The receptor’s conformation -and consequently, its activity-is modulated by post-translational modifications and interactions with ligands, cofactors, and chromatin [6]. GR’s quaternary structure appears essential for its function, but controversy remains regarding which oligomeric state (monomer, dimer, tetramer) might be responsible for triggering specific gene programs [7].

Unliganded GR is primarily cytoplasmic and thus unable to access the genome. To be able to bind its ligand, GR’s LBD requires the aid of a complex chaperone cycle, including Hsp90, Hsp70, the adaptor proteins p23, and the immunophilins FKBP51 and FKBP52 [8-11]. Upon glucocorticoid binding, the receptor undergoes a conformational change that facilitates its translocation to the nucleus, where it regulates the expression of hundreds of genes [1, 6].

How GR moves from the cytoplasm to the nucleus is still unresolved. First, the receptor should reach the perinuclear region and, once there, it must interact with a nuclear pore complex (NPC) [12] and pass through it. While there are several proposed mechanisms for each of the stages involved in the nuclear import [13], two apparent mutually exclusive models prevailed. The classical model proposed that ligand binding in the cytoplasm triggers the release of GR from the chaperone heterocomplex, exposing the otherwise occluded nuclear localization signals (NLS) located in the hinge region and LBD. This free or *transformed* GR can be subsequently recognized by the importin system, leading to an importin-mediated nuclear traffic [13]. An alternative model gives a more prominent role to dynein-mediated microtubule transport, actively moving GR towards the nuclear periphery [14]. Interestingly, under this model, GR is not released from the chaperone heterocomplex, wherein the receptor interacts indirectly with dynein through FKBP52. It has even been proposed that the entire heterocomplex traverses the nuclear pore complex, and it is only in the nucleoplasm that GR is finally released [15].

Another controversial issue is whether GR dimerizes before, during, or after passing through the NPC and into the nucleus. As GR is complexed with chaperones and accessory proteins in the absence of ligand, all structural data points to only monomers passing through the pore, as both DBD and LBD, known to participate in GR-GR dimer formation [16-18], would be sterically impaired to interact with each other [8]. However, some reports claimed to have measured dimeric unliganded GR in the cytoplasm [19-21], but offer no structural explanation as to how the dimer can form if GR is still coupled to the chaperone heterocomplex. Similarly, the oligomeric state of liganded GR during its journey towards the nucleus is also controversial, as there have been reports claiming that GR travels either as a monomer [20, 22] or as a dimer [23]. In any case, once in the nucleus, it appears that most of the free, diffusive GR is dimeric [16, 17, 24], although there have been reports of a mixed population of monomers and dimers [21, 25]. The oligomeric state of GR bound to chromatin is less understood and harder to measure experimentally, but cumulative evidence points to DNA-dependent tetramerization of GR [17, 18, 26, 27].

Here, we explore the mechanisms governing GR nuclear translocation and the relevance of its quaternary structure in this process by using the pair correlation function (pCF) approach. This method quantifies the correlation between fluorescence intensity fluctuations observed at two different positions within a sample and produced by single fluorescent molecules moving from one position to the other [28]. This method has been previously used to measure the molecular flow of tracer fluorescent proteins through the NPC [29], among other applications [30].

We show that both GR monomers and dimers are able to translocate into the nucleus, demonstrating that dimerization can precede nuclear import. Interestingly, while GR monomers seem to approach the NPC’s cytoplasmic side by passive diffusion, GR dimers also use the faster microtubule-dynein active transport system to reach the pores efficiently. In fact, the perinuclear vimentin intermediate network contributes to faster transport, likely by constraining GR molecules near microtubules. Therefore, our work resolves apparently contradicting reports on the quaternary structure of GR during nuclear import but reopens the door for a microtubule-dependent, heterocomplex-independent GR translocation.

## Methods

### Cell culture and sample preparation for imaging

D4 mouse adenocarcinoma cell lines were described previously [26, 31]. Cells were cultured in Dulbecco’s modified Eagle’s medium (DMEM, Gibco) supplemented with 10% fetal bovine serum (FBS, Internegocios S.A.) and 1% penicillin-streptomycin (Gibco). To prevent expression of the integrated tet-off system, 5 μg/mL tetracycline (Santa Cruz Biotechnology) was added to the culture medium. Cells were grown at 37^◦^C under a humidified atmosphere with 5% CO_2_.

For microscopy measurements, cells were grown overnight on sterilized 25-mm round coverslips placed into 35-mm plates with 1.0 ml of complete medium. This medium was replaced with DMEM containing 5% charcoal-stripped FBS 24 h before observation.

D4 cells expressing Halo-tagged proteins were incubated for 1 h with 50 nM JF646 (Janelia Farms, HHMI, USA, [32] and then washed twice for 5 min before imaging. Hormonal stimulation was performed by incubating the cells with 100 nM dexamethasone (DEX) (Sigma-Aldrich) or 10 μM 21-hydroxy-6,19-epoxyprogesterone (21OH-6,19OP, [33]). To promote the partial depolymerization of the microtubule network, the cells were incubated with 16 μM nocodazole for 1 h. The procedures described previously were performed at 37°C and 5% CO_2_.

### Plasmids and transfection

Cells grown on coverslips were transiently transfected using Lipofectamine 2000 (Invitrogen) according to the vendor’s instructions and observed 24 h after transfection. Briefly, ~2.10^5^ cells were cultured on coverslips as described above, transfected with 1– 1.5 μg of DNA, and the transfection medium was replaced with DMEM containing 5% charcoal-stripped FBS. Cells were incubated overnight in this medium before microscopy experiments.

The plasmids were HP1α-GFP (Addgene #17652); EMTB-3xGFP [34], which codifies the microtubule-binding domain of ensconsin fused to three tandem copies of GFP (Addgene # 26741), kindly provided by Dr. Arpita Upadhyaya (University of Maryland, College Park, MD). The dominant-negative construct containing the head and alpha-helical domain 1A of vimentin GFP-vim(1-138) [35] and GFP-p50 were gifts from Dr. Vladimir Gelfand (Northwestern University, Chicago, IL); pNup153-EGFP (Euroscarf P30650); pHalo-GRmon [36] and pEGFP-GR [37] were previously described.

### Microscopy

Confocal images were acquired in a Zeiss LSM 980 confocal microscope using a Plan-Apochromat 63x oil immersion objective (NA= 1.4). EGFP or GFP, and JF646 were excited with solid-state diode lasers of 488 nm and 639 nm, respectively. The average power at the sample was ~1 μW (488 nm) and ~5 μW (639 nm). The fluorescence emitted by the sample was detected using photomultipliers set in the photon-counting detection mode and configured to register in the range between 490-552 nm (EGFP or GFP) and 641-694 nm (JF646).

Time-lapse images were acquired in the absence of ligand and every 5 min during hormonal stimulation over 30 min. Images were registered with pixel size and dwell time of 132 nm and 1.02 μs, respectively.

Line-scanning experiments in the presence of ligands were run between 5-30 min from hormone addition. In these experiments, the lasers were set to repetitively scan along a line (pixel size=105 nm, line length =13.4 μm) across the nuclear envelope, simultaneously. The line acquisition frequency and total number or lines were 1961 Hz and 2.10^5^, respectively. To avoid photodamage, we run a single line-scanning experiment per cell.

Microscopy measurements were performed at 37ºC and 5% CO_2_.

### pCF analysis

The two-color line-scanning data were used to generate kymographs for the channels corresponding to GR (GR-JF646; GRmon-JF646 or EGFP-GR) and to protein labels used to delimit the cytoplasm-nucleus boundary (HP1α-GFP, GFP-p50, GFP-vim(1-138) or Nup153-EGFP). The data showing evident motion of the nucleus were excluded from further analysis.

We selected regions 315 nm-wide (3 pixels) in the GR kymographs equidistant to the nuclear boundary (defined in the second kymograph) and separated from each other by 1.05 μm (10 pixels). For each subcellular region, we computed the autocorrelation function (ACF) as:

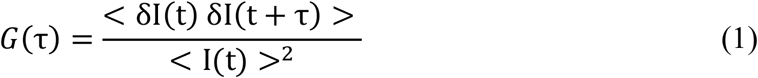

where I(t) is the fluorescence intensity at time t, τ is the lag time, the brackets indicate average values over the time-course of the experiment, and δI (t)=I(t)−<I(t)> represents the fluorescence intensity fluctuation at time t. We discarded the files in which the ACF data was zero.

To estimate G_0_ at the nuclear region, the ACF data were fitted with the following equation, accounting for anomalous diffusion [38]:

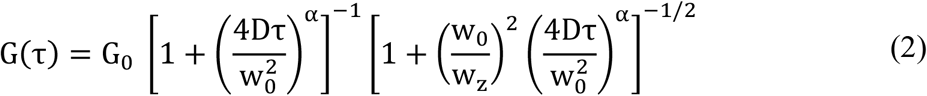

where D is the diffusion coefficient, w_0_ and w_z_ are the radial and axial waist of the point spread function, α is the anomalous diffusion exponent. G_0_ = γ/N, where N is the mean number of molecules in the confocal volume and γ is a geometric factor that depends on the detection profile and corresponds to 0.35 for a confocal setup [39].

We did not apply the diffusion-plus-two-binding model used in our previous work [40], as the reduced number of data points and the lower temporal resolution of the line-scanning experiments compared to single-point FCS measurements lead to increased noise in the ACF data.

The pair correlation function (pCF) was calculated as [28]:

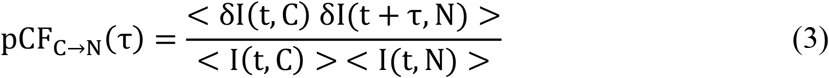

where I(t, C) and I(t, N) are the fluorescence intensity at time t registered at the cytoplasmic (C) and nuclear (N) regions.

In most of the experiments, each pCF_C→N_ curve was normalized by the value of G_0_ at the nucleus obtained from the fitting of its nuclear ACF (Eq. 2). This relative pCF_C→N_ is related to the proportion of GR molecules detected at the cytoplasmic position that are observed afterwards in the nucleus, following a similar reasoning as described previously [41]. In the conditions where the nuclear intensity was very low (i.e., experiments in the absence of ligand), we did not calculate the relative pCF as ACF data were too noisy.

We excluded from the analysis pCF or cross-pCF data (see below) with values equal to zero across the entire lag-time range, as these indicate that no translocation events were detected. Such cases may arise, for example, from the absence of functional nuclear pores within the laser-scanned region.

FCS and pCF analysis were performed using the SimFCS program (LFD, Irvine, CA, USA). The mean pCF or cross-pCF curves (see below) were smoothed using an adjacent-averaging method (20-point window) in Origin software.

### CCF and cross-pCF analyses

The two-color cross-correlation function (CCF) at each subcellular region was calculated as follows [42]:

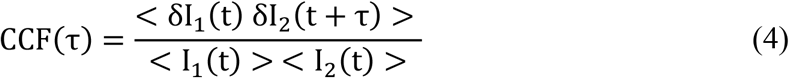

where I_1_ (t) and I_2_ (t) are the fluorescence intensity at time t registered in channels 1 and 2, respectively. The CCF curve was fitted with Eq. 2 to facilitate data visualization.

The two-color pCF function was calculated as follows [28]:

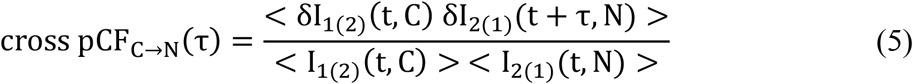

where I_1(2)_ (t, C) and I_1(2)_ (t, N) are the fluorescence intensity at time t registered at channels 1(2) at the cytoplasmic (C) and nuclear (N) regions.

CCF and cross-pCF analysis were performed using the SimFCS program (LFD, Irvine, CA, USA).

### Quantification of GR nuclear import

The nucleus area was delimited from whole cell confocal images using the transmission images as a reference (see, for example, **Supplementary Fig. 1A**), and the mean nuclear intensity of GR-JF646 and GRmon-JF646 at a single cell level was quantified using the Fiji software.

For each nucleus, the mean nuclear intensity (I) registered as a function of the incubation time (t) was fitted with Eq. 6 to obtain the maximum value reached (I_plateau_) (see, for example, **Supplementary Fig. 1B**). Then, the intensity obtained at every incubation time was normalized by I_plateau_.

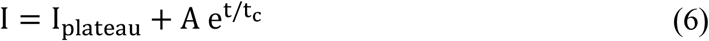

where A is a constant, and t_c_ is the characteristic import time.

The GR import level (I_30_/I_0_) was calculated at a single cell level as the ratio between the normalized intensity values obtained at 30 min and in the control condition.

### Statistical analysis

The values of the intensity ratio (I_30_/I_0_) were expressed as median ± standard error. To verify whether the medians (med) of data groups (g) were significantly different, we performed a hypothesis test computing the p-values as follows [43]:

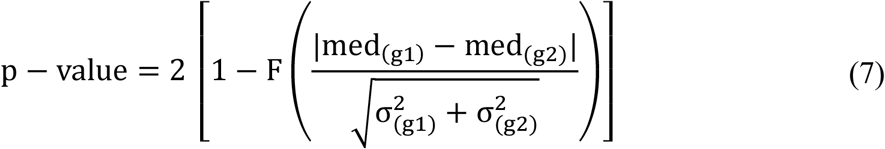

where F is the standard normal distribution and σ^2^ _(g1)_ and σ^2^ _(g2)_ are the variance of each data set. Differences were regarded as significant at p < 0.05.

The variance and the median standard error were computed by a bootstrap procedure [44]. Statistical data analysis was performed using R software.

## Results

### Visualizing GR translocation from the cytoplasm to the nucleus by the pair correlation function (pCF) analysis

We used the D4-HaloGR murine mammary adenocarcinoma cell line, wherein endogenous GR has been knocked out [26, 31]. These cells stably express HaloTag-GR at endogenous levels [36] and were conjugated with the HaloTag ligand JF646 [45] for live cell fluorescent microscopy. Stimulation with dexamethasone (DEX), a potent glucocorticoid agonist, triggers the translocation of GR bound to JF646 (GR-JF646) into the nucleus, reaching a steady state after ~25-30 min of incubation **(Fig. 1A-B and Supplementary Fig. 1**). These imaging experiments provide ensemble and spatially averaged information and, therefore, they do not reveal the underlying molecular mechanism(s) driving this process. To dissect these mechanisms, we used the pCF approach [28] that can provide information on the journey of single molecules between two positions within a cell.

**Fig. 1.**
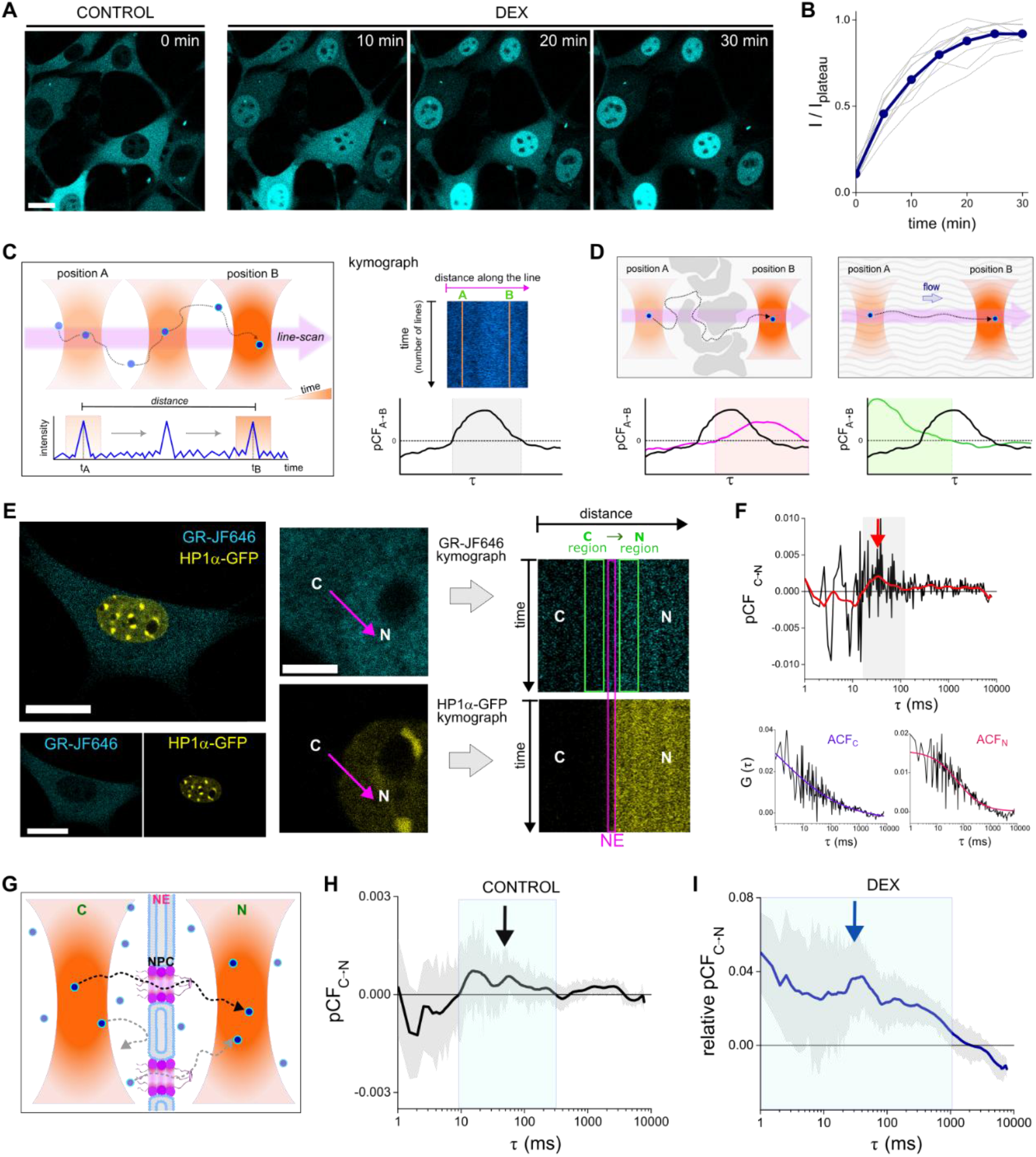
Nuclear translocation of GR unveiled by pCF analysis. **(A)** Representative confocal images of D4-HaloGR cells labeled with JF646 before (CONTROL) and during dexamethasone (DEX) stimulation. Scale bar: 20 µm. **(B)** The fluorescent intensity curves obtained for each cell (gray) were processed as described in Methods, and the average data (dark blue) is represented as a function of time. (N_cells_= 9). **(C)** Schematic illustration of the line-scan experiment and pCF analysis. (left panel) The microscope is set to repetitively scan the sample along a line (pink arrow), registering the fluorescence emitted by the molecules (blue circles) diffusing through the confocal volume (orange). (right panel) The intensity collected along the line, as a function of time, is used to construct a kymograph and calculate the pCF function (bottom panel) between two positions selected by the user (A, B). The positive correlation region (gray rectangle) corresponds to the characteristic time (τ_c_) required for molecules to move from position A to B. **(D)** Cartoon representing pCF data expected when obstacles delay molecules diffusion (left panel; pink line) or with an active flow (right panel; green line). **(E)** (left panel) Representative confocal image of a D4-HaloGR cell (cyan, GR-JF646) expressing HP1α-GFP (yellow, nucleus) in control condition. Scale bar: 20 µm. (right panel) Zoom-in region of the cell after 10 min of DEX stimulation. Scale bar: 5 µm. In pCF experiments, the laser was set to scan a line repetitively (arrow) from the cytoplasm (C) to the nucleus (N). HP1α kymograph was used to determine the position of the nuclear envelope (NE) and define the cytoplasmic and nuclear regions (green rectangles) selected in the GR-JF646 kymograph for pCF analysis. **(F)** pCF and ACF curves (black) obtained in representative experiments. (top panel) Smoothed pCF curve (red). The arrow points to the characteristic positive correlation region. (bottom panel) ACF curves corresponding to cytoplasmic (ACF_C_) and nuclear (ACF_N_) regions were fitted with Eq. 2 to facilitate data visualization. **(G)** Cartoon illustrating the pCF analysis applied to study GR translocation. Due to the dimension of the confocal volume (orange), pCF analysis allows not only the study of GR translocation through the nuclear pore complex (NPC), but also its dynamics near the nuclear envelope (NE) at the cytoplasmic (C) and nuclear region (N). For simplicity, we did not include in the scheme other cell components that may be involved in GR translocation. **(H)** pCF curve obtained for GR-JF646 in the absence of ligand (CONTROL). pCF data obtained for each cell was used to compute the mean pCF curve and then smoothed (black) as described in Methods. The standard error is shown in gray. The arrow points to the characteristic positive correlation region. (N_cells_=17). **(I)** Relative pCF curve obtained for GR-JF646 during dexamethasone (DEX) stimulation over 30 min. pCF data obtained for each cell was normalized by their respective G_N_(0) and then averaged (see Methods). The plot shows the smoothed mean pCF curve (blue) and the standard error (gray). The arrow points to the characteristic positive correlation region. (N_cells_=37).

Briefly, in a confocal microscope, the laser is set to scan relatively fast (~millisecond range) and repetitively a line in the sample that intersects the positions of interest (points A and B in **Fig. 1C**), and the fluorescence intensity is collected pixel-by-pixel. In the simplest case, where fluorescent molecules passively diffuse within the sample, some of the molecules detected in position A will subsequently reach position B. As molecules produce intensity fluctuations when diffusing through the confocal volume, each molecule moving from A to B will produce a fluctuation in the intensity trace collected at both positions but separated by a characteristic time lag (τ_c_). The τ_c_ value depends on the diffusion coefficient (i.e., how fast molecules move) and the distance between the selected positions.

These correlated fluctuations can be detected by calculating the pCF function (Eq. 3, see Methods), which provides information on the number of molecules moving from positions A to B as well as the value of τ_c_. **Figure 1D** schematize more complex scenarios in which obstacles delay the transit of molecules from positions A to B, producing a delayed pCF peak or, conversely, an active flow pushes molecules from position A to B, shifting τ_c_ to lower values. These examples illustrate how pCF analysis could be a useful approach to extract detailed insights into the journey of molecules between different positions of interest with minimal perturbation.

To study the molecular mechanism(s) of GR translocation, we run pCF experiments in D4-HaloGR cells labeled with JF646. To visualize the cell nucleus during the pCF experiments, we either transiently expressed nuclear proteins such as heterochromatin protein HP1α-GFP (**Fig. 1E**) or Nup153-EGFP to also locate the nuclear envelope (NE) (**Supplementary Fig. 2A**). Interestingly, Nup153-EGFP labeling allowed us to determine that positions A and B ought to be at least 525 nm (i.e., 5 pixels) away from the nuclear envelope to ensure they are indeed located in the cytoplasm and the nucleus, respectively (**Supplementary Fig. 2B**).

The lasers were set to repetitively scan a line that intersected the nuclear envelope (**Fig. 1E**). Then, the data were represented as kymographs and used to calculate the pCF function between the intensity traces collected at the selected positions in the cytoplasm and the nucleus, herein referred to as pCF_C→N_ (see Methods) (**Fig. 1F**). Additionally, we calculated the autocorrelation function (ACF) at both positions and determined G_N_(0) from the fitting at the nuclear position (**Fig. 1F**, see also Methods).

Considering the dimensions of the confocal observation volume (radial and axial radii of ~200 nm and ~1 μm, respectively [46], the pCF_C→N_ data will contain information of the mechanism(s) involved in both the searching process of GR molecules nearby the nuclear pore complex (NPC) combined with the passage through the pore itself (**Fig. 1G**). Throughout this work, the term *translocation* will encompass the exploratory motion of GR within a region nearby to the nuclear pore complex, and the transit through the pore into the nucleus. In addition, we will use the term *nuclear import* to refer to the process involving the motion of GR from the whole cytoplasm into the nucleus which is usually observed by ensemble-average methods such as whole-cell imaging.

In the absence of ligand, the pCF_C→N_ function shows a positive region with local maxima at 15-50 ms (**Fig. 1H**, arrow), in the range of the characteristic time reported for the translocation of Imp α-JF549 (~14 ms, [47]) and slightly higher than that of the smaller mCherry-NLS (~3.5 ms, [48]); the latter determined by orbital tracking and pCF. These authors also showed that mCherry, without any NLS thus expected to passively diffuse through the pore [49], presents a broader distribution of longer transit times up to 0.5-1 s [48]. Therefore, the relatively narrower distribution observed for the transit times of unliganded GR molecules suggests that the importin-dependent system is required for their translocation, as expected given the relatively large size of the GR–HaloTag fusion protein (120–130 kDa).

In contrast, the pCF_C→N_ data in DEX-stimulated cells showed a wider positive correlation region that spans the 1-1000 ms range with a local peak at ~30 ms (**Fig. 1I**, arrow), suggesting that additional or other mechanism(s) account for the C→N flow of liganded GR-JF646 molecules.

We also run pCF experiments in cells stimulated with the rigid steroid 21-OH-6,19OP [50], as the receptor bound to this ligand presents incomplete nuclear import (**Supplementary Fig. 3A-B**), [16]. The pCF_C→N_ data of GR-21-OH-6,19OP showed a delayed translocation with respect to GR–DEX, as no positive correlation is detected in the 1-20 ms range (compare **Supplementary Fig. 3C with Fig. 1I**), suggesting that the translocation of the receptor, when bound to 21OH-6,19OP, occurs over a longer time window.

Taken together, our results thus far demonstrate the feasibility of applying the pCF approach to study GR translocation and highlight the dependence of the translocation on ligand-dependent GR conformation.

### Microtubules-dynein favor GR translocation

As mentioned above, previous whole-cell imaging experiments demonstrated a role of the microtubule network in aiding GR import [51], likely by providing the tracks for the active transport of GR from the cytoplasm towards the perinuclear zone. However, whether they are also required for completing the translocation process remains controversial [52]. In addition, growing evidence points to direct and indirect connections between microtubules and NPCs that could contribute to precisely locating cargoes at NPCs [53]. To get further insights into the molecular mechanisms involved in the translocation of active GR, we run pCF experiments under conditions wherein the microtubule network is compromised.

First, D4-HaloGR cells labeled with JF646 were incubated with nocodazole (1 h at 37°C), inducing a partial depolymerization of microtubules (**Fig. 2A**). Whole-cell imaging experiments after DEX-stimulation show that this treatment did not abolish GR import (**Fig. 2B**) but reduced this process **(Fig 2C**) in line with previous observations [51, 54].

**Fig. 2.**
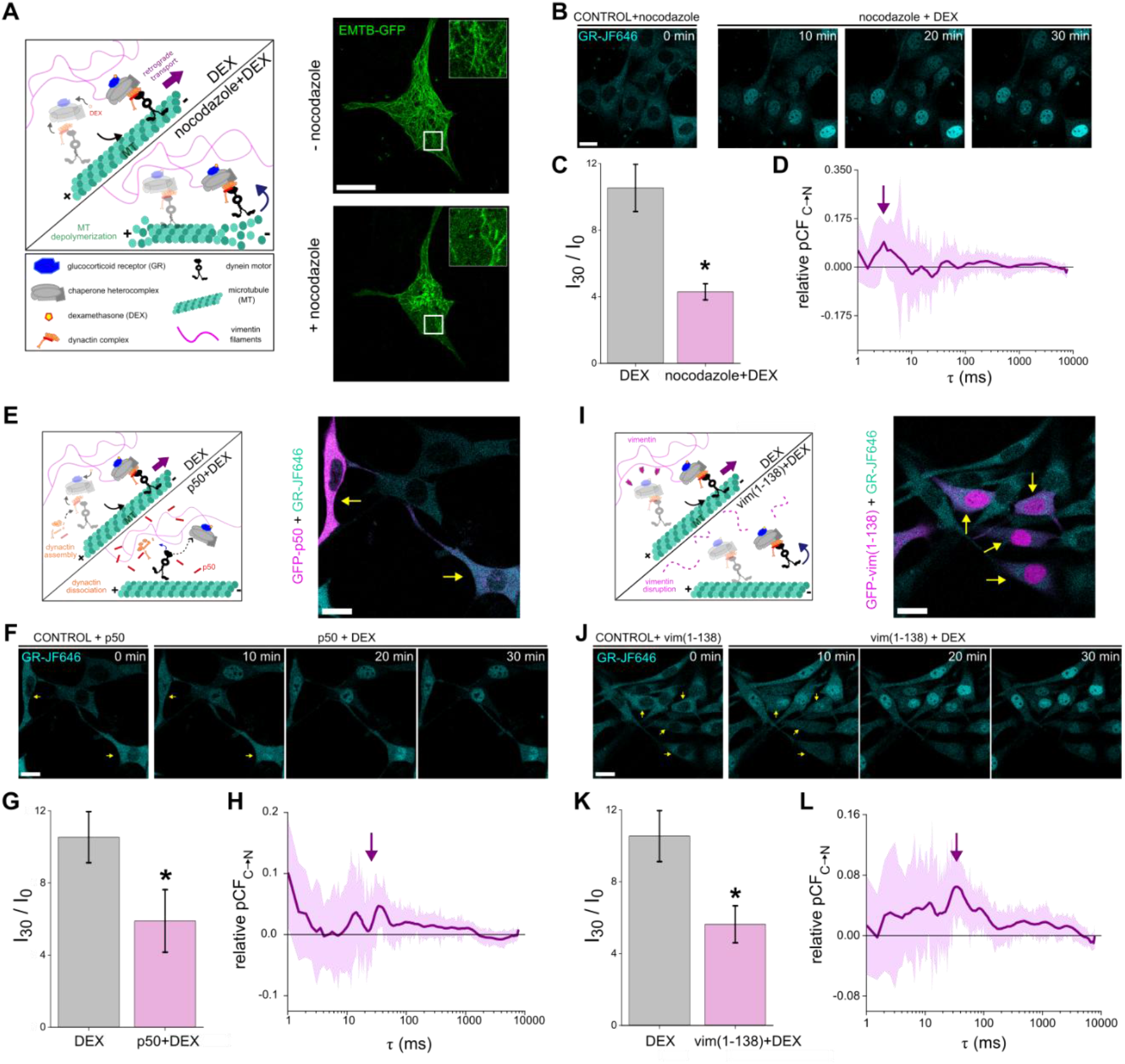
Liganded GR translocation requires dynein-mediated transport along microtubules. **(A)** (left panel) Cartoon showing the retrograde transport model for GR-heterocomplex driven by dynein along microtubules (MT) upon dexamethasone stimulation before (DEX) and after nocodazole treatment (nocodazole+DEX). (right panel) Representative confocal image of a D4-HaloGR cell transfected with EMTB-3xGFP to label microtubules (green) registered before and after incubation with nocodazole. Inset: Zoom-in region indicated by the rectangle. Scale bar: 20 µm. **(B)** Confocal images of D4-HaloGR cells labeled with JF646 (cyan, GR-JF646) and treated with nocodazole registered in the absence of ligand (CONTROL) and at different incubation times with DEX. Scale bar: 20 µm. **(C)** Nuclear import level of GR-JF646 in the presence of DEX in untreated and nocodazole-treated cells. The data is shown as median ± standard error. (N_cells-DEX_= 9 and N_cells-nocodazole_= 12). * p < 0.05. **(D)** Relative pCF curve obtained for GR-JF646 during DEX stimulation in nocodazole-treated cells. The plot shows the smoothed mean pCF curve (purple) and standard error (light purple). The arrow points to the positive correlation region. (N_cells_=16). **(E)** (left panel) Cartoon illustrating the action of dynactin p50 subunit overexpression on GR-heterocomplex transport. (right panel) Representative confocal image of D4-HaloGR cells expressing GFP-p50 (magenta, arrows) and labeled with JF646 (cyan, GR-JF646) registered in the absence of ligand (CONTROL). Scale bar: 20 µm. **(F-G)** Comparison of the nuclear import level of GR-JF646 in untreated cells and those expressing GFP-p50 (arrows). The data is shown as median ± standard error. (N_cells-DEX_= 9 and N_cells-p50_= 10). * p < 0.05. **(H)** Relative pCF curve obtained for GR-JF646 during DEX stimulation in GFP-p50 transfected cells. The plot shows the smoothed mean pCF curve (purple) and the standard error (light purple). The arrow points to the positive correlation region. (N_cells_= 16). **(I)** (left panel) Cartoon showing the role of vimentin filaments in preventing GR-dynein complex detachment from the MT during transport. (right panel) Representative confocal image of D4-HaloGR cells expressing GFP-vim(1-138) (magenta, arrows) and labeled with JF646 (cyan, GR-JF646) registered in the absence of ligand (CONTROL). Scale bar: 20 µm. **(J-K)** Comparison of the nuclear import level obtained for GR-JF646 in untreated cells and those expressing GFP-vim(1-138). The data is shown as median ± standard error. (N_cells-DEX_= 9 and N_cells-vim(1-138)_ = 11). * p < 0.05. **(L)** Relative pCF curve obtained for GR-JF646 during DEX stimulation over 30 min in GFP-vim(1-138) transfected cells. The plot shows the smoothed mean pCF curve (purple) and the standard error (light purple). The arrow points to the characteristic positive correlation region. (N_cells_= 15).

We next ran pCF experiments in nocodazole-treated cells after DEX-stimulation and observed a qualitative change in the shape of the normalized pCF_C→N_ curve. Specifically, the wide range of positive correlation is lost and the pCF_C→N_ curve exhibits a positive region with local maxima at ~3 ms **(Fig. 2D**, arrow). Thus, disrupting the microtubule network affects GR translocation dynamics. These results support that active transport along microtubules probably constitutes a faster and more efficient mechanism than passive diffusion to concentrate GR molecules from the peripheral to the perinuclear regions. Nevertheless, we cannot rule out confounding effects arising from this harsh treatment, and therefore, we followed alternative approaches to gain further insights into the role of microtubules in GR translocation.

As dynein transports GR molecules from the cell periphery to the perinuclear region [55, 56], we hypothesize that it may also help position and anchor the receptor near, or even at, the nuclear pores. This localization would increase the likelihood of their interaction, an event that would likely occur less efficiently if GR were to rely solely on passive diffusion within the perinuclear region.

To test this hypothesis, we disrupted dynein-microtubule interactions by overexpressing p50 fused to GFP (**Fig. 2E**). p50 is a subunit of the dynactin complex [57] which interacts with dynein, allowing the binding of the motor to its cargoes and also increasing its processivity [58]. Overexpression of p50 leads to the disassembly of dynactin [59], effectively preventing the interaction of GR with the dynein motor [60].

Consistent with previous reports [56], imaging experiments revealed an impaired incorporation into the nucleus of DEX-stimulated GR-JF646 in D4-HaloGR cells overexpressing the p50 dynactin subunit (**Fig. 2F-G**). Under these conditions, pCF assays reveal changes in the translocation dynamics (**Fig. 2H**). In contrast to the wide positive correlation region observed in control cells, the normalized pCF_C→N_ curve rapidly decreases and presents a local maximum at ~ 12 and 35 ms (**Fig. 2H**, arrow). The change in the shape of the pCF_C→N_ curve observed in the ~1-10 ms region after p50 overexpression (compare **Fig. 1I vs Fig. 2H**) suggests that microtubules and dynein motors are involved in the rapid search and nuclear pore translocation of GR molecules occurring within this time window. The positive correlation observed at τ < 5 ms could arise from the small fraction of assembled dynactin complexes that still form, as their inhibition depends on p50 expression levels.

Finally, we evaluated the role of intermediate vimentin filaments in GR translocation as this cytoskeleton network contributes to maintaining motor-cargo complexes close to microtubules, preventing their detachment [61-63]. We expressed the dominant-negative construct vim(1-138) fused to GFP in D4HaloGR cells to disrupt the vimentin network [35] and analyzed GR translocation after DEX stimulation in those cells expressing the mutant vimentin (**Fig. 2I**). Imaging experiments show that disruption of vimentin filaments impaired GR nuclear import (**Fig. 2J-K**). The pCF_C→N_ curve revealed a slightly lower contribution in the 1-3 ms (compare **Fig. 1I vs Fig. 2L**), suggesting that the vimentin network contributes to the fast GR transport also consistent with the proposed role of this network in maintaining cargoes close to microtubules.

These results suggest that perinuclear microtubules via dynein motors keep active GR molecules close to NPCs, an interaction that seems to be favored by the vimentin network and may be part of a faster mechanism for GR translocation to the nucleus.

### GR dimers and monomers translocate into the nucleus through different mechanisms

There have been contradictory reports on when and where GR dimerizes, either after ligand activation in the cytoplasm, during translocation [23], or only in the nuclear compartment [15]. Therefore, we used the pCF method to elucidate whether GR monomers and/or dimers can translocate into the nucleus and the mechanism(s) involved in their translocation.

We first studied the translocation of GRmon, a monomeric GR that combines mutations in both the DBD and the LBD dimerization surfaces (GRA465T/I634A, [16]). Imaging experiments performed in D4 cells co-expressing HaloGRmon (JF646) and HP1α-GFP show an impaired nuclear import of GRmon-JF646, in comparison with GR (**Fig. 3A-B**).

**Fig. 3.**
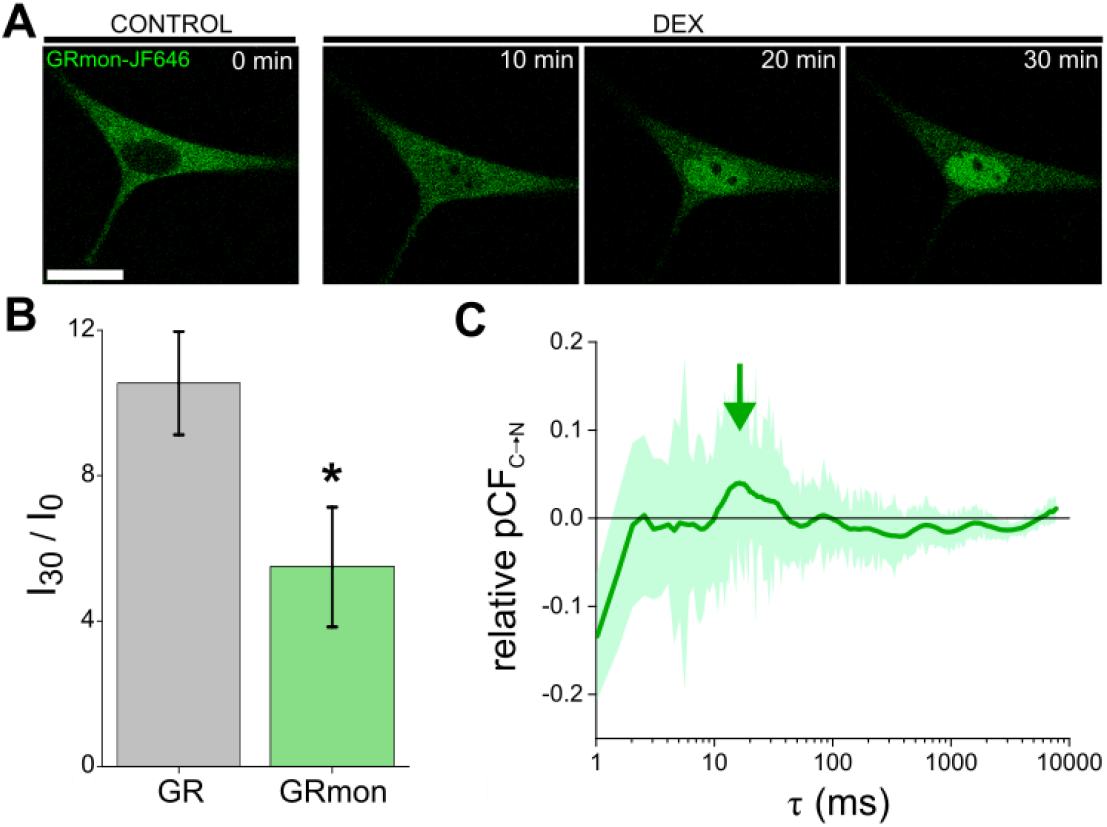
Monomeric GR exhibits slower nuclear translocation. **(A)** Representative confocal images of a D4 cell expressing HaloGRmon labeled with JF646 (green, GRmon-JF646) registered in the absence of ligand (CONTROL) and during DEX stimulation. Scale bar: 20 µm. **(B)** Comparison of the nuclear import level obtained for GRmon-JF646 (GRmon) and GR-JF646 (GR). The data is shown as median ± standard error. (N_cells-GRmon_= 7 and N_cells-GR_= 9). * p < 0.05. **(C)** Relative pCF curve obtained for GRmon-JF646 during dexamethasone (DEX) stimulation. The plot shows the smoothed mean pCF curve (green) and the standard error (light green). The arrow points to the characteristic positive correlation region. (N_cells_=10).

Next, we run pCF experiments in cells expressing GRmon-JF646 and observed that the normalized pCF_C→N_ curve presented a single peak at τ ~15-20 ms (**Fig. 3C**), indicating a slower translocation than that observed for the wild type receptor. Notably, the τ_c_ value determined for GRmon-JF646 lies in a similar time-range to that observed for unliganded GR-JF646 (compare **Fig. 1H with Fig. 3C**). Notably, the absence of a positive correlation at lower τ values (compare **Fig. 1I to 3C**) suggests GRmon search process near the nuclear pore relies solely on passive diffusion, without involvement of the microtubule–dynein system.

To explore whether GR can translocate into the nucleus as a dimer, we run two-color pCF experiments [28] in D4-HaloGR cells labeled with JF646 and transiently expressing EGFP-GR. In these experiments, the lasers scan a line crossing the nucleus repetitively, and the intensities of the EGFP- and JF646-labeled GR molecules are collected simultaneously into two independent detectors. We first calculated the two-color cross-correlation function [64] at single positions selected from the kymographs near the nuclear envelope (Eq. 4 in Methods, **Fig. 4A-B**). The positive cross-correlation suggests the presence of GR dimers in the nucleus and perinuclear (cytoplasmic) regions.

**Fig. 4.**
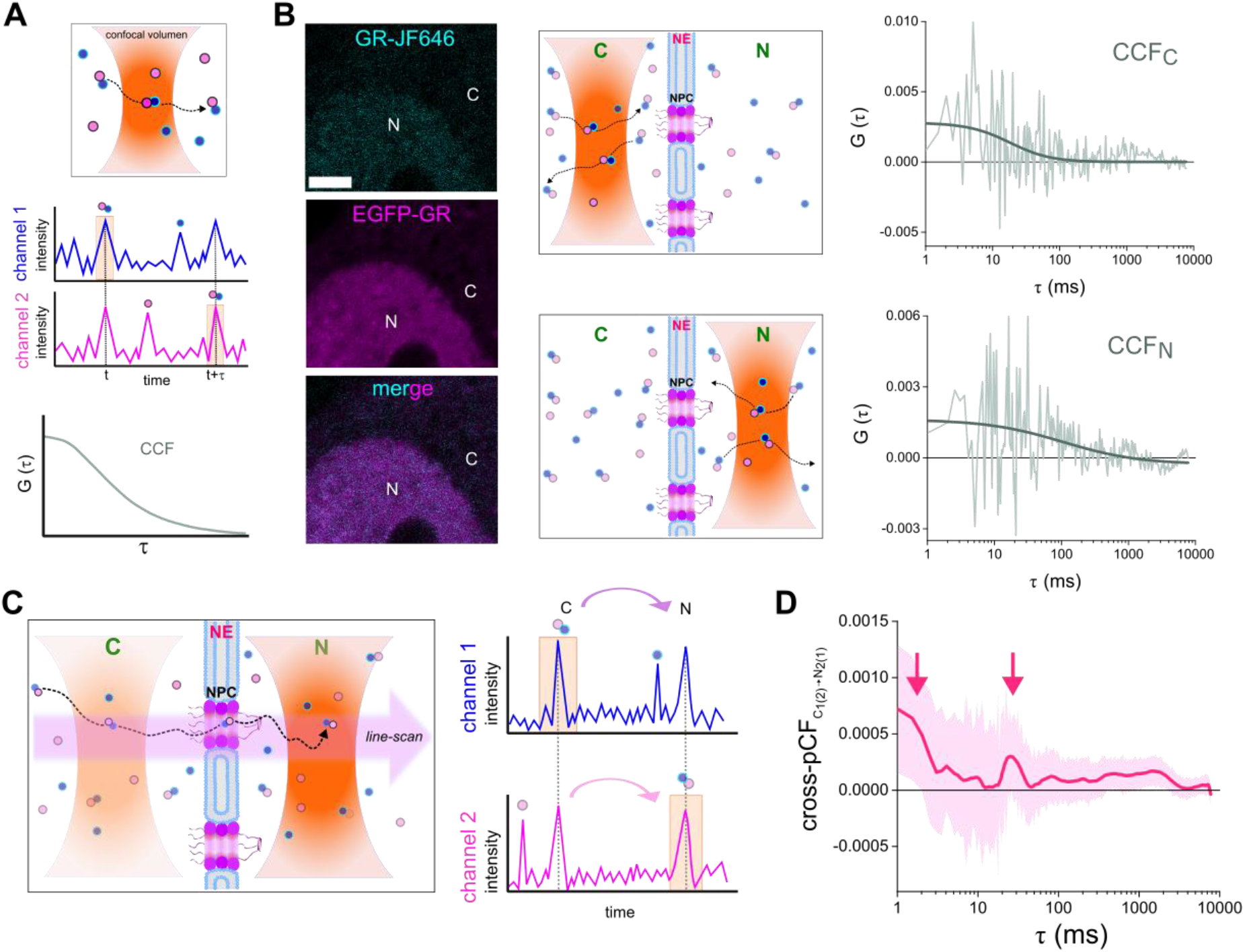
A GR subpopulation translocates as a dimer. **(A)** Schematic illustration of the CCF analysis. The fluorescence emitted by the molecules (blue and magenta circles) diffusing through the confocal volume (orange) are registered at two independent detectors according to their color (channel 1 and channel 2). The intensity profiles obtained for each channel are processed to compare the fluorescence fluctuations over time, computing the CCF (Eq. 4). The positive cross-correlation curve supports the presence of two-color dimers. **(B)** CCF analysis. (left panel) Representative confocal images of a D4-HaloGR cell labeled with JF646 (cyan, GR-JF646) and expressing EGFP-GR (magenta) during stimulation with DEX. C, cytoplasm; N, nucleus. Scale bar: 3 µm. (middle panel) Scheme illustrating GR monomers (single color) and dimers (two color) near the nuclear envelope (NE), moving also through the confocal volume (orange). The CCF analysis was performed at the cytoplasmic (CCF_C_) and nuclear (CCF_N_) regions. (right panel) Representative CCF curves obtained at each cellular compartment; the data were fitted with Eq. 2 to facilitate their visualization. **(C)** Schematic illustration of the two-color pCF analysis. The microscope is set to repetitively scan the laser on the sample along a line (pink arrow), and the fluorescence emitted by molecules (blue and magenta circles) moving through the confocal volume (orange) in two independent detectors (channel 1 and 2). The intensity trajectories obtained in the channels provide information on two-color dimers moving from the cytoplasm to the nucleus. **(D)** Cross-pCF curve obtained for GR-JF646/EGFP-GR in DEX-stimulated cells. The plot shows the smoothed mean cross-pCF curve (pink) and the standard error (light pink). The arrows point to the positive correlation regions. (N_data_=40 and N_cells_=20).

Then, we calculated the cross-pair correlation function (cross-pCF_C→N,_ Eq. 5) between the cytoplasmic and nuclear positions (**Fig. 4C**) defined from the kymographs as described before. This data provides information on the journey of two-color complexes moving from one position to the other.

The cross-pCF curve showed two positive regions at τ ~1-3 ms and 20-40 ms (**Fig. 4D**, arrows), indicating that GR-JF646 and EGFP-GR can indeed translocate together as dimers. Since the higher τ_c_ value was similar to the positive correlation region determined when the dynein complex was impaired (**Fig. 2H**), it is tempting to speculate that this slower mechanism involves GR dimers moving toward the pore through passive diffusion. The two peaks observed in **Fig. 2H** could, in this context, reflect the monomers and dimers diffusing near the NPC and passing through it. However, additional experiments will be required to test this hypothesis. In the same line of reasoning, the fast correlation appears to correspond to the cytoskeleton-driven transport of GR dimers. In fact, the absence of a positive pCF correlation in this time lag range for GRmon (**Fig. 3C**) suggests that monomeric GR is unable to move by active transport.

In conclusion, our data indicate that both dimers and monomers can translocate into the nucleus, but through different strategies: GR dimers can move through both cytoskeleton-dependent transport and passive diffusion, while GR monomers rely solely on passive diffusion.

## Discussion

The precise localization of proteins within the eukaryotic cell is essential for life, and thus, it is tightly regulated [65]. Transport in and out of the nucleus occurs mainly through NPCs that enable a rapid and selective transit of a wide variety of biomolecules. While small molecules below a soft threshold of ~30-60-kDa [49] can passively diffuse through NPCs, larger proteins require the energy-dependent importin system to translocate into the nucleus.

Although a rough comparison of GR monomeric size (94-97 KDa [1]) with this threshold suggests the necessity of active translocation, exactly what happens after ligand binding remains disputed [14]. Some aspects of this process have been studied in whole-cell imaging experiments, but unfortunately, these methodologies average out the behavior of large populations of unsynchronized molecules and thus relevant aspects of the mechanism(s) of translocation remain hidden. For example, whether GR translocates as a dimer or a monomer, and whether each species uses different mechanisms to reach the NPC, remains controversial. Even though the nuclear import and export of GR was employed as a proof of principle to validate methods such us massively parallel FCS [66, 67] or multipoint holographic FCS [68], the mechanisms behind GR translocation were not explored.

In recent years, advanced fluorescence microscopy techniques have revolutionized the study of biological processes by enabling the exploration of single biomolecules moving within living cells [69]. These techniques are minimally invasive and provide exquisite information on the mechanisms of molecular processes *in situ*. The last years showed seminal contributions on NPC function, which is now considered not only as a molecular sieve but also playing an active role in the unidirectional transport of molecules based on their size, its mechanical properties, and charge distribution [49, 70, 71]. For example, a recent study showed that cargoes with low mechanical stability regions adapt better to pore restrictions and show higher translocation efficiencies [71]. 3D single-molecule tracking [72] and 3D MINFLUX [73] revealed that cargoes move preferentially through the pore walls instead of using the central cavity. MINFLUX data also indicate that the trajectories of importing and exporting molecules overlap on the pore surface. Unfortunately, these studies were run after permeabilizing the cells with digitonin, a treatment that, while minimally affecting the pore function, still removes several components of the cytoplasm, including those related to cytoskeletal-related transport [72, 74]; thus preventing them from observing how molecules reach the NPC from the cytoplasmic side. Finally, 3D orbital tracking of the NPC combined with pCF analysis of fluctuations produced by fluorescent cargoes moving through the pore revealed a single characteristic nuclear import time in the millisecond range, whereas passively diffusing small molecules present a broad distribution of longer import times [48].

In this context, the pCF approach emerged as a sophisticated tool to study the movement of single biomolecules between different cellular regions and, particularly, between the cytoplasm and the nucleus in intact cells. Thus, we applied this methodology to explore the nuclear translocation of GR. Our data gives a unique perspective on the quantitative aspects of GR translocation. Specifically, we found that the inactive receptor can enter the nucleus with characteristic translocation times of ~15-50 ms, close to that observed for the translocation of the Imp α-JF549 (14 ms, [47]) and mCherry-NLS with the combination of 3D tracking with pCF [48](~3.5 ms). The higher translocation time determined for GR-JF646 could be due to differences in molecular properties, including its size, and/or the shorter distance between the cytoplasm and nuclear positions in that work (240 nm, [48]) compared to ours (525 nm).

As mentioned above, GR is a relatively large molecule, and it is also expected to be associated with a chaperone heterocomplex to maintain its inhibited conformation and ligand binding capabilities [75]. Therefore, our results suggest that a small fraction of unliganded GR molecules may expose the NLS, reach the NPC by diffusion, and translocate into the nucleus. The low GR nuclear concentration in unstimulated cells suggests either an efficient nuclear export of unliganded GR molecules or a fast degradation of nuclear GR in the absence of glucocorticoids. Unfortunately, the low concentration of GR in the nucleus makes it difficult to study the mechanism of unliganded GR export by pCF analysis. Moreover, recent data also suggest degradation as the most likely scenario [76].

On the other hand, our pCF analysis revealed that the characteristic times of the translocation of DEX-stimulated GR span a wider range, indicating that additional mechanisms, apart from diffusion, produce a rapid translocation of the active receptor. Relevantly, the ligand can modulate this process, as pCF analysis in the presence of the synthetic ligand 21OH-6,19OP shows a delayed translocation of the receptor with respect to DEX-stimulated GR, suggesting that GR conformation affects translocation pathways.

It has been postulated that GR uses the microtubule network to efficiently reach the perinuclear region [14, 75]. In this context, we evaluated the contribution of microtubules and dynein motors in the translocation process. Our data support that microtubules provide tracks to efficiently transport DEX-stimulated GR to the proximity of nuclear pores via interaction with dynein motors through dynactin. Relevantly, we also verified that the vimentin intermediate filaments network contributes to this process, probably by preventing the dynein/dynactin/GR complexes from detaching from the tracks.

Finally, we evaluated the oligomeric state of GR while translocating into the nucleus. Our pCF and cross-pCF analysis demonstrates that both monomers and dimers can translocate to the nucleus. However, in a forced monomeric conformation (GRmon), translocation is detected only between ~15-50 ms, in the range of the translocation time interval measured when the dynactin complex is disrupted. Thus, monomeric GR appears to approach the NPC only through diffusion. In contrast, the cross-pCF detecting GR dimers revealed that dimers show two discrete translocation characteristic times of ~1-3 and 20-40 ms that probably represent GR molecules making use of microtubule-dependent transport or passive diffusion to approach the NPC from the cytoplasm side, respectively.

We should highlight the overall complexity of the GR translocation process, which involves GR monomers and dimers, a combination of passive diffusion and active transport, as well as the influence of the local microenvironment of the NPCs (e.g., the density of the vimentin network and microtubules). Moreover, in our pCF experiments, the laser crosses the nuclear envelope at random (blind) positions, and thus, the number of pores included in the analyzed region may vary between experiments. This likely contributes to the observed variability among pCF curves. In fact, all these factors contribute to local variations in the translocation process and, consequently, to broader distributions of translocation times, as observed, for example, for DEX-stimulated GR.

Taken together, our data suggest that 1) dimerization occurs in the cytoplasm after ligand activation, 2) both monomers and dimers can translocate into the nucleus, and 3) GR monomers cannot use the microtubule network efficiently. Therefore, it is tempting to speculate that the translocation time distribution observed in DEX-stimulated cells results from both monomers and dimers entering the nucleus.

These results challenge a few aspects of the current model for GR translocation, specifically whether GR travels bound to the chaperone heterocomplex through the microtubule network [15]. Since GR dimerization [18, 27, 77] appears sterically incompatible with the receptor also being bound to this heterocomplex [8, 9], it is tempting to propose that GR dimers must be travelling dissociated from chaperones. This will also predict a direct interaction of GR with FKBP52-dynein, which has been reported elsewhere [55]. However, how or whether GR dimers can interact with FKBP52 remains an open question, as a recent structural study not only confirms a direct GR-FKBP52 interaction, but also implies that dimerization and FKPB52 binding are sterically incompatible [8]. On the other hand, monomeric GR, while sterically able to use the microtubule network through the chaperone-FKBP52-dynein complex, appears very inefficient at it. Overall, our data are compatible with both chaperone-dependent and independent microtubule-directed and diffusive pathways in the control of the nuclear localization of GR.

## Declaration of competing interests

The authors declare that they have no competing interests.

## Funding sources

This research was partially supported by ANPCyT (PICT 2020-00818 to V.L., PICT PRH 2018-0573 to D.M.P.), Universidad de Buenos Aires (UBACyT 20020190100101BA to V.L.) but PICT funding disbursements were suspended two years ago, following the Argentina government’s decision to cut financial support for scientific research across the country.

## Author contribution

**María Cecilia De Rossi:** Conceptualization, Methodology, Investigation, Formal analysis, Data Curation, Validation, Visualization, Writing - Original Draft, Writing - Review & Editing, Project administration.

**Diego Martín Presman:** Conceptualization, Methodology, Validation, Visualization, Resources, Writing - Original Draft, Writing - Review & Editing, Funding acquisition, Project administration.

**Valeria Levi:** Conceptualization, Methodology, Validation, Visualization, Writing - Original Draft, Writing - Review & Editing, Funding acquisition, Project administration, Supervision.

## Notes

### Competing Interest Statement

The authors have declared no competing interest.

