## Supplemental Figures 1-3 for "Pair correlation function analysis revealed different nuclear translocation mechanisms for glucocorticoid receptor’s monomeric and dimeric forms"

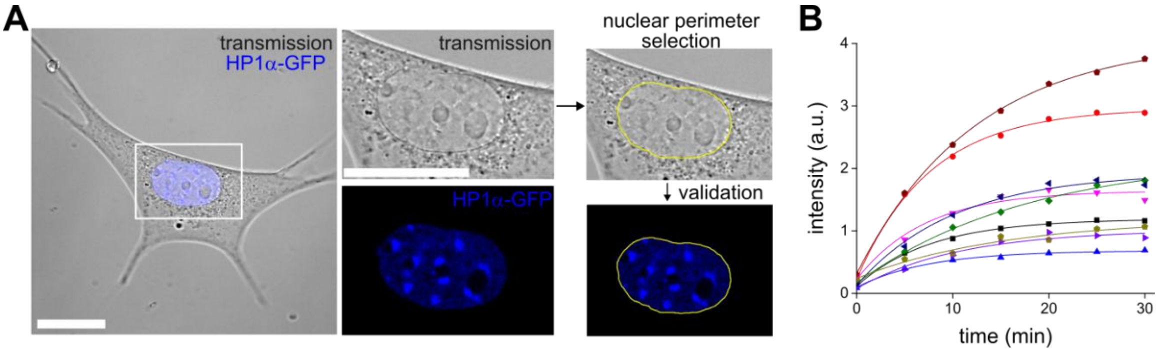


**Supplementary Fig.1. Quantification of GR nuclear intensity. (A)** Determination of the nuclear area. An example of how the nuclear area was delimited using transmission and confocal images. Scale bar: 20 µm. **(B)** Representative examples of the nuclear intensity (in this case, of GR-JF646) registered as a function of the incubation time after DEX addition. Each data set correspond to a different cell and was fitted using Eq.6 (continuous lines).

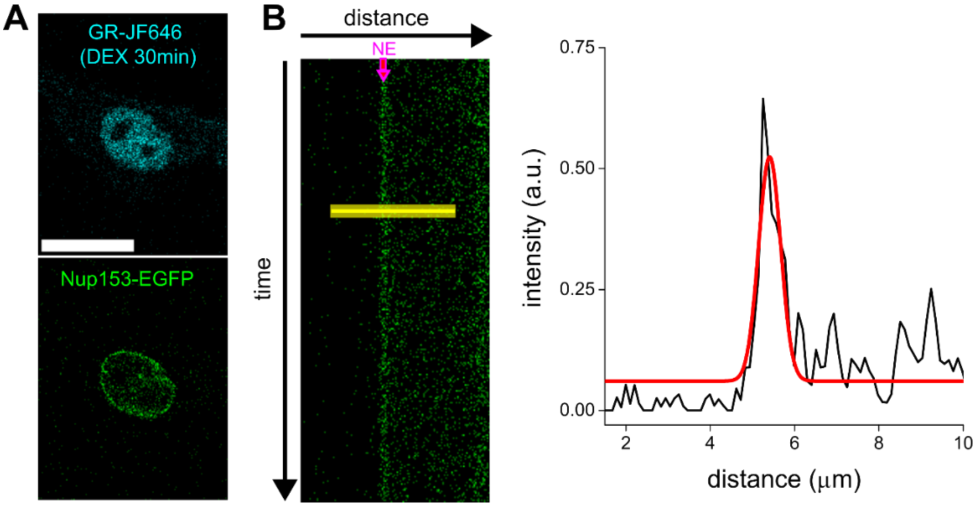

**Supplementary Fig.2. Determination of the distance between the cytoplasmic and nuclear regions for pCF analysis. (A)** Representative confocal images of a D4-HaloGR cell labeled with JF646 and transfected with Nup153-EGFP. Scale bar: 20 µm. **(B)** Determination of the width of the nuclear envelope (NE) image. (left panel) Kymograph obtained for Nup153-EGFP. (right panel) The fluorescent intensity profile (black) registered along the yellow line was fitted with a Gaussian function (red) to estimate the NE width (FWHM= 0.62 ± 0.06 µm).

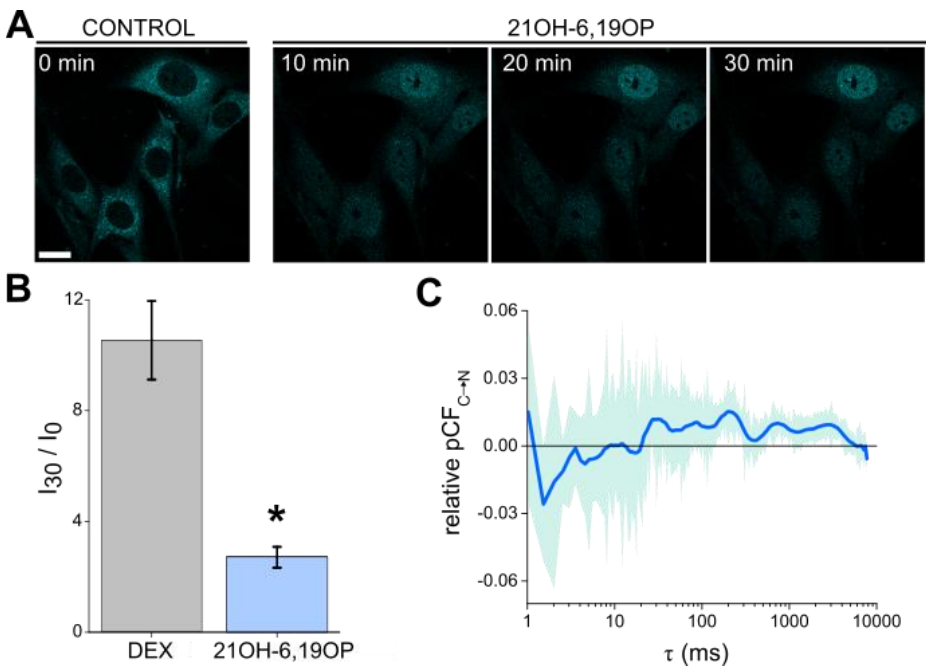


**Supplementary Fig.3. Translocation of GR bound to 21OH-6,19OP. (A)** Representative confocal images of D4-HaloGR cells labeled with JF646 before and during 21OH-6,19OP stimulation. Scale bar: 20 µm. **(B)** Comparison of the nuclear import level for GR-JF646 stimulated with DEX or 21OH-6,19OP. The data is shown as median ± standard error. ($N_{cells\text{-}DEX}$= 9 and $N_{cells\text{-}21OH\text{-}6,19OP}$= 12). * $p < 0.05$. **(C)** Relative pCF curve obtained for GR-JF646 during 21OH-6,19OP stimulation. The plot shows the smoothed mean pCF curve (blue line) and the standard error (light blue). ($N_{cells}$=9).
